# Signal-to-noise ratio of event-related fields in on-scalp and off-scalp MEG

**DOI:** 10.64898/2026.08.17.744953

**Authors:** Mainak Jas, Teppei Matsubara, Abbas Sohrabpour, Padmavathi Sundaram, Maria Mody, Seppo P. Ahlfors

## Abstract

Optically pumped magnetometer (OPM) sensors can be placed closer to the scalp than conventional superconducting quantum interference devices (SQUID), resulting in larger magnetoencephalography (MEG) signals from neuronal activity. For event-related sensor data, such as epileptogenic activity or sensory and motor evoked responses, however, OPMs and SQUIDs often differ less in signal-to-noise ratio (SNR) than in signal magnitude. We examined two factors contributing to the relative SNR: the dependence of the signal magnitude on source depth and the effect of scalp-to-sensor distance on the noise level. Simulated MEG data for a current dipole in a spherical head model confirmed that on-scalp sensor placement delivers the largest SNR gain for superficial sources. Depending on the relative overall noise level, there may be a crossover source depth at which SNR is equal for on-scalp and off-scalp sensors and beyond which off-scalp sensors achieve higher SNR. Analysis of the equal-SNR source depth in different-sized spherical head models indicated that, for a given relative noise level, the proportion of the brain where SNR is higher in OPM than in SQUID was larger in small head models, supporting the benefits of OPMs in pediatric studies. To experimentally evaluate noise contributions of brain and non-brain origin to the SNR, we recorded somatosensory evoked fields (SEFs) at varying scalp-to-sensor distances. Generally, both the evoked response magnitude and the noise level were lower when the sensors were further away from the scalp; consequently, the SNR depended less than the signal magnitude on the scalp-to-sensor distance. Comparison of power spectral densities (PSDs) at different sensor-to-scalp distances allowed us to identify whether the dominant noise source was of brain or non-brain origin at different frequency bands. Overall, the results highlight complementary properties of OPMs vs. SQUIDs in terms of SNR, which is of interest when optimizing MEG experiments for specific subject populations and brain regions.

## 1. INTRODUCTION

In magnetoencephalography (MEG), electrical activity in the human brain is observed non-invasively by recording magnetic fields outside the head (Ahlfors & Mody, 2019; Cohen, 1972; Supek & Aine, 2019). In general, it is desirable to place the MEG sensors as close as possible to the head, because of the dependence of the signal magnitude on the distance between the neural source and the sensor (Cuffin & Cohen, 1979; Gaetz, Otsubo, & Pang, 2008; Tanaka, Ahlfors, & Stufflebeam, 2025; Tripp, 1983). MEG instruments commonly have several hundreds of Superconducting Quantum Interference Device (SQUID) sensors, which provide high-quality low-noise MEG data for many clinical and research applications (Vrba & Robinson, 2001). Typically, fixed helmet-shaped arrays of SQUID sensors are immersed in liquid helium inside a cryogenic Dewar vessel. The thickness of the Dewar wall sets a lower limit on the scalp-to-sensor distance. The minimum scalp-to-sensor distance in conventional SQUID systems is about 20 mm. With advanced SQUID-based instrumentation, such as high-Tc SQUIDs or cryocooled sensors, however, this distance can be substantially reduced, enabling even near “on-scalp” MEG recordings (Andersen et al., 2017; Okada et al., 2016; T. P. Roberts et al., 2014). Optically Pumped Magnetometers (OPM), have recently emerged as an alternative to SQUIDs for MEG (Boto et al., 2018; Budker & Romalis, 2007). OPMs do not require cryogenic cooling, therefore allowing on-scalp placement. Other on-scalp technologies currently under development include NV-diamond- and semiconductor-based sensors (Barry et al., 2020).

Shorter scalp-to-sensor distance is expected to improve the spatial resolution of neural source localization (Iivanainen, Stenroos, & Parkkonen, 2017; Nugent, Benitez Andonegui, Holroyd, & Robinson, 2022; Wens, 2023). Signal magnitudes in OPM sensors are higher than in off-scalp SQUID sensors by a factor of about 4–8 (Brickwedde et al., 2024). For event-related data, such as epileptic discharges or sensory evoked responses, however, OPMs and off-scalp sensors differ less in SNR than in signal magnitude (Borna et al., 2020; Boto et al., 2017; Feys et al., 2022; Iivanainen, Zetter, & Parkkonen, 2020; Marhl, Jodko-Wladzinska, Bruhl, Sander, & Jazbinsek, 2022; Wang et al., 2024). In the present study, we examined the signal and noise properties of on-scalp and off-scalp MEG sensors to better understand factors contributing to the SNR.

MEG signals are influenced by several kinds of noise, including intrinsic sensor noise, external environmental noise, and participant-related noise (Taulu, Simola, Nenonen, & Parkkonen, 2019). Intrinsic sensor noise is commonly of the order of 2-5 fT/√Hz for SQUIDs and 7-30 fT/√Hz for OPMs (Brookes et al., 2022) and uncorrelated between sensors. Environmental noise consists of magnetic fields from external sources, such as nearby moving objects (e.g., trucks, elevators, or chairs) or electrical equipment. Another source of noise in MEG is participant-related magnetic fields, either of physiological origin (e.g., ocular or cardiac), or due to magnetized materials on the body (e.g., dental or other implants). Both external and subject-related artefacts are usually highly correlated across MEG sensors and therefore can be effectively reduced by signal processing techniques (Seymour et al., 2022; Taulu et al., 2019).

For event-related activity, after pre-processing to reduce artefacts and external interferences, the dominant source of noise in MEG data is often ongoing background brain activity (M. Hämäläinen, Hari, Ilmoniemi, Knuutila, & Lounasmaa, 1993). This “brain noise” can be defined as magnetic fields due to brain activity that is unrelated to the activity of interest in a given study. Brain noise is correlated across MEG sensors, and the spatial patterns (scalp topographies) can be similar to those of the event-related activity. An effective way to reduce brain noise is to average over repeated events, although this may not always be feasible, e.g., for infrequent epileptic discharges or real-time brain-computer interface applications.

Detection of deep sources has traditionally been challenging for MEG because signals from deep sources are much weaker than signals from superficial sources (Cuffin & Cohen, 1979; Goldenholz et al., 2009; Hillebrand & Barnes, 2002). Given that on-scalp sensors are closer to the deep structures, one might expect that on-scalp sensors provide an advantage over off-scalp sensors for detecting deep sources. However, simulations have indicated that the highest relative SNR for on-scalp compared with off-scalp sensors occurs for superficial sources (Boto et al., 2017; Iivanainen et al., 2017; Matsubara et al., 2026). The benefit of on-scalp sensors for superficial sources can be partially attributed to the strong nonlinear dependence of the signal on the source-to-sensor distance. Another potentially contributing factor is that not only the signal magnitude but also the noise level can depend on the scalp-to-sensor distance. Since on-scalp sensors are closer than off-scalp sensors to the brain, the data are expected to contain more brain noise.

In the present study we examined the influence of the source depth and the scalp-to-sensor distance on the SNR in MEG. First, we simulated SNRs as a function of source depth at different relative levels of noise in on- and off-scalp sensors. The simulations suggested that there may be an equal-SNR source depth, at which on-scalp and off-scalp sensors have the same SNR. For sources located deeper than the equal-SNR depth, off-scalp sensors yielded higher SNR than on-scalp sensors. Second, we recorded somatosensory evoked magnetic fields (SEFs) with OPM and SQUID sensors placed at varying distances from the scalp. Altering the scalp-to-sensor distance allowed us to infer whether the dominant noise was of brain or non-brain origin, aiding the interpretation of the simulated and experimental SNR values in OPMs and SQUIDs. Overall, our goal was to clarify complementary properties of OPMs and SQUIDs, which is of interest to researchers devising strategies to optimize the SNR in MEG for specific subject populations and brain regions.

## 2. METHODS

### 2.1. Simulations of SNR as a function of source depth

We simulated the SNR in on-scalp and off-scalp sensors as a function of the location (depth) of a source of interest as well as the relative level of noise between the sensor types. The source was assumed to be a current dipole, which is a widely used source model for many types of event-related MEG signals (Hari et al., 2018; Scherg & Von Cramon, 1986).

For convenience, we use the terms “OPM” and “SQUID” here interchangeably with “on-scalp” and “off-scalp” sensors, respectively. It should be noted, however, that SQUID sensors made of high-Tc superconductors can be practically on-scalp (Andersen et al., 2017), whereas OPM sensors typically have non-zero scalp-to-sensor distance (Knappe, Sander, & Thrams, 2019). Currently, OPMs are the most practical near-on-scalp sensors; however, NV-diamond- and semiconductor-based sensors are currently under development (Barry et al., 2020). Our simulations are expected to yield practical guidelines for target SNR levels in any types of sensors.

Simulated MEG signals were calculated for a current dipole located at different depths in a spherical head model, which allows analytical expressions for the magnetic field (M. Hämäläinen et al., 1993; Sarvas, 1987; Tripp, 1983). The simulation geometry is illustrated in **Fig. 1**. The tangentially oriented dipole moment vector was **Q** = *Q***e**_y_, where the dipole magnitude *Q* was chosen to be 30 nAm; e_y_is the unit vector in the *y*-direction. The dipole was located at **r**_Q_ = *r*_Q_e_z_, where *r*_Q_ is the distance from the sphere origin along the *z*-axis. The value of *r*_Q_ was varied between 0 and *b*, the radius of the brain. The radius of the head (scalp surface) was denoted by ℎ. We chose several different values for *b* and ℎ (**Table 1**), representing head sizes from newborn to adult (Beauchamp et al., 2011; Guo, Roche, & Moore, 1988; Lew, Hämäläinen, Ahlfors, & Okada, 2021; Zahran et al., 2022).

**Figure 1:**
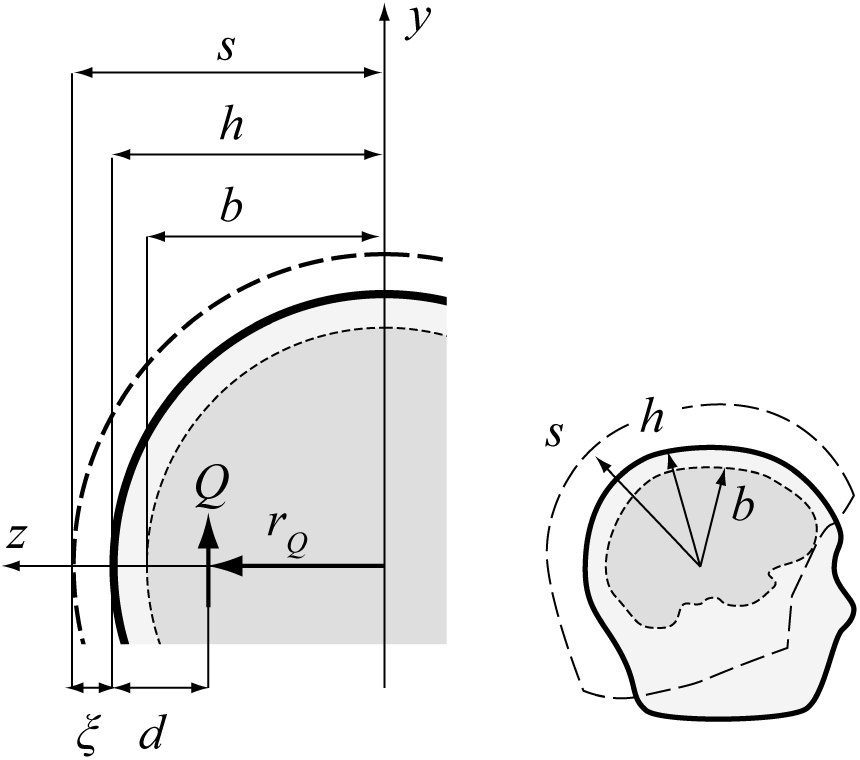
Geometry of the spherical head model used in the simulations. A tangential dipole ***Q*** was assumed to be at a distance *r*_Q_ from the sphere origin. The radii of the brain and scalp (head) were *b* and ℎ, respectively. Thus, the depth of the source measured from the scalp surface was *d* = ℎ − *r*_Q_. The sensors were assumed to be on a spherical surface with radius *s*; the scalp-to-sensor distance was ξ = *s* − ℎ. The inset at right illustrates *b*, ℎ, and *s* in a schematic side view of the head.

**Table 1:** Head model parameters (in mm) used in simulations.

| Head Model | Head radius ($h$) | Brain radius ($b$) | $(h - b)^*$ |
| --- | --- | --- | --- |
| Infant, newborn | 55 | 48 | 7 |
| Infant, 1 year | 70 | 62 | 8 |
| Child, 8 years | 85 | 73 | 12 |
| Adult | 95 | 80 | 15 |

\*Combined thickness of CSF, skull, and scalp

All sensors were assumed to measure the radial field component *B*_r_ = **B** · **e**_r_ of the magnetic flux density **B**. SQUID sensors were on a spherical surface with the radius *s* = ℎ + ξ, where ξ is the scalp-to-sensor distance. We used ξ = 18 mm, which corresponds to the minimum distance between the sensors and the scalp in the Triux neo SQUID system (MEGIN, Finland). The OPM sensors were assumed to be located on-scalp, i.e., at distance ℎ from the origin (ξ = 0). In the simulations, we did not consider the finite size of the SQUID pick-up coils or the OPM sensor cells over which the magnetic field is integrated in practical devices.

The radial component of the magnetic field at location **r**_s_ for a current dipole is (M. Hämäläinen et al., 1993; Tripp, 1983)

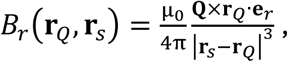

The spatial topography of *B*_r_ has peaks symmetrically on each side of the dipole **Q**, in our case in the *xz*-plane. Note that in the sphere model, the magnetic field does not depend on ℎ or *b* (Ilmoniemi & Sarvas, 2019). For sensors in the *xz*-plane at the distance *s* from the origin, the sensor position is

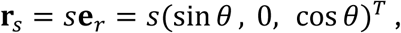

where *θ* ∈ [0, π] theta is the polar angle between **r**_s_ and the *z*-axis. Setting ∂*B*_r_/∂*θ* = 0 yields for the peak of the topography the angle θ_0_:

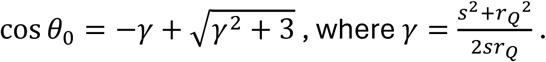

The corresponding maximum value of *B*_r_ is (Boto et al., 2017):

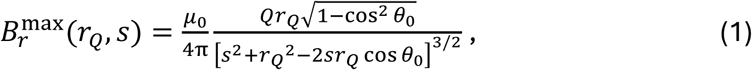

which was used as the signal magnitude in the simulations.

Denoting the noise power for the two sensor types as *σ*^2^_OPM_ and *σ*^2^_SQUID_, the SNR values were defined as:

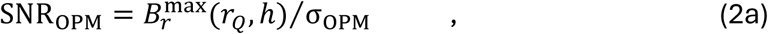

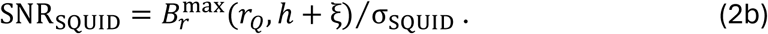

For mathematical convenience, we defined the source location here in terms of the radius *r*_Q_ (distance from origin). In the Results section, however, we used the more intuitive “source depth”, i.e., the distance between the source and the scalp surface. In the sphere model, the source depth is simply *d* = ℎ − *r*_Q_, and thus, in Eq. (2), ℎ − *d* can replace *r*_Q_.

To compare the SNR between OPM and SQUID sensors, we varied the relative noise level

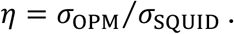

For reasons elaborated in the Discussion section, we assumed that *η* ≥ 1, i.e., the noise was never lower in OPMs than in SQUIDs. The dipole location for which SNR_OPM_ = SNR_SQUID_ for a given value of *η* was defined as the *equal-SNR source depth* (*d*_eq_). If *d*_eq_ exists within the brain (i.e., in the range ℎ − *b* ≤ *d*_eq_ < ℎ), it can be solved by equating Eqs. (2a) and (2b), yielding the implicit equation for *d*_eq_:

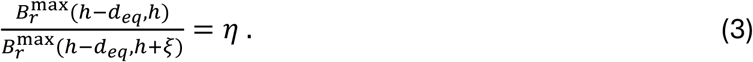

In the simulations, we parametrically varied the dipole depth *d* (from deep to superficial), the relative noise level *η* (from 1 to 6), and the head size (from infant to adult, **Table 1**). In addition, to relate the simulated data to our SEF recording, we varied the scalp-to-sensor distance ξ (from on-scalp to 6 cm off-scalp).

### 2.2. Somatosensory evoked fields at different scalp-to-sensor distances

To evaluate how the signal magnitude and SNR vary as a function of the scalp-to-sensor distance, we recorded somatosensory evoked fields (SEFs) using multiple different placements of OPM and SQUID sensor arrays. In contrast to the simulations, where the source depth was varied, for the real data we varied the sensor locations but assumed that the source location was constant. i.e., the source of the N20 response to median nerve stimulation. We did not attempt to systematically vary the source depth in this experiment.

SEFs were recorded in a 62-year-old healthy male participant. The study protocol was approved by MGH Institutional Review Board, and informed consent was obtained. The subject was seated upright, instructed to fixate on a marker, keep eyes open, and minimize head movements throughout the experiment. The median nerve in the right wrist was electrically stimulated using a constant current stimulator with 2-ms pulses, the amplitude of which adjusted to elicit a visible thumb twitch. The stimulus onset asynchrony was 500 ms. Each run consisted of 500 stimulus trials.

OPM data were recorded with 11 single-axis magnetometer sensors (FieldLine Gen2, FieldLine, Boulder, CO) in a magnetically shielded room with a pair of aluminum and mu-metal layers (Imedco AG, Haegendorf, Switzerland). The anti-alias low-pass filter cut-off was 330 Hz and sampling rate was 1000 Hz. The sensors were mounted on a 3D-printed helmet, fitted with precision probe holders and adjustable spacers to control the scalp-to-sensor distance (**Fig. 2A**). The center of the OPM sensors was ∼7 mm from the inside surface of the helmet in the “on-scalp” position (cubical vapor cell with 10-mm side length on a 2-mm-thick helmet shell). The sensors were measuring the magnetic field component normal to the scalp surface.

**Figure 2:**
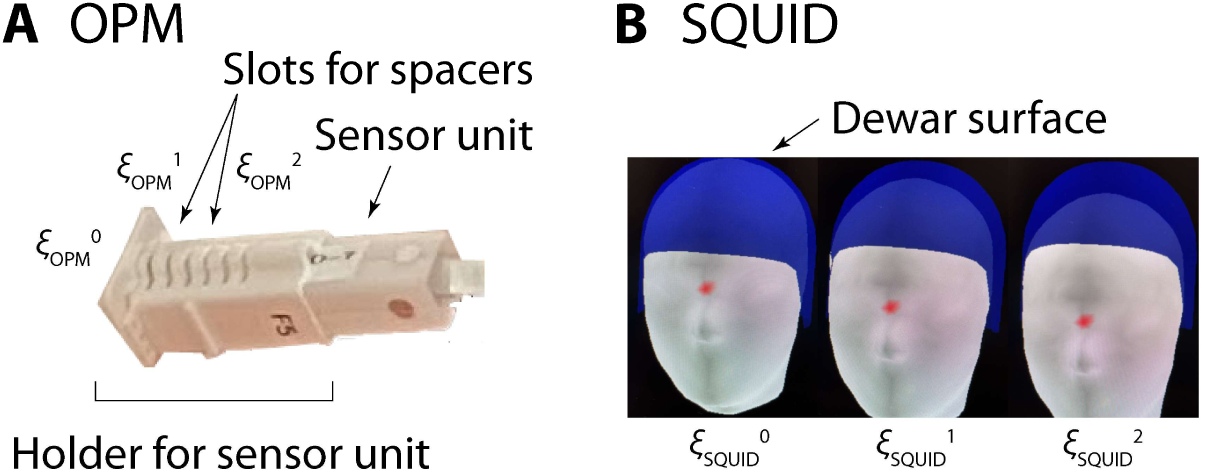
Measuring MEG signals at varying distances from the scalp. **A**: OPM sensors were attached to a 3D-printed helmet with holders for each sensor unit. Each sensor unit holder had several precision-made openings, 5 mm apart from each other, for removable spacers, which allowed controlling the scalp-to-sensor distance of the OPM sensors. Here, no spacers are depicted, as the sensor unit is at the closest position to the scalp. **B**: SQUID sensors were in a rigid array inside a liquid helium Dewar. The scalp-to-sensor distances were varied by having the participant’s head slide down inside the helmet-shape Dewar.

SQUID data were recorded using a 306-channel Triux neo system (MEGIN, Helsinki, Finland) with 102 magnetometers, measuring the field component approximately normal to the scalp, and 204 planar gradiometers, inside a magnetically shielded room with 3 pairs of aluminum and mu-metal layers (Imedco) (Cohen, Schläpfer, Ahlfors, Hämäläinen, & Halgren, 2002). Only magnetometer data were used here, to facilitate comparison with the OPMs. Anti-alias low-pass filter cut-off was 330 Hz and sampling rate was 1000 Hz. The thickness of the wall of the helmet-shaped Dewar was 18 mm, setting a lower limit for the scalp-to-sensor distances for the SQUID sensors. The head position with respect to the SQUID sensor array was determined using 5 Head Position Indicator (HPI) coils on the scalp.

OPM and SQUID data were collected in separate sessions, each with 4 runs. In the first run, the sensors were as close to the scalp as possible, (ξ^0^_OPM_, ξ^0^_SQUID_). In the second and third runs, the scalp-to-sensor distance was increased by about 1 and 2 cm, respectively, from the initial position, (ξ^1^_OPM_ and ξ^2^_OPM_, ξ^1^_SQUID_ and ξ^2^_SQUID_). The fourth run was a replication of the first, with sensors again close to the scalp (ξ^0,rep^_OPM_, ξ^0,rep^_SQUID_). In addition, empty room data was collected without the participant.

For the OPMs, the sensor position was adjusted by placing spacers in the 3D-printed probe holders, increasing the scalp-to-sensor distance by 10 or 20 mm (**Fig. 2A**). For SQUIDs, the scalp-to-sensor distance for sensors over the somatosensory cortex was varied by lowering the participant’s seat such that the head slid down within the helmet (**Fig. 2B**); the relative position of the head and the SQUID sensor array was determined using the HPI coils; the overall shifts from the initial head position (ξ^0^_SQUID_) were 10 mm (ξ^1^_SQUID_), 23 mm (ξ^2^_SQUID_), and 3 mm (ξ^0,rep^_SQUID_). Since the SQUID sensors were in a fixed array inside the Dewar, the temporal and occipital sensors could not be moved away from the scalp; therefore, we focused on the central (frontoparietal) sensors near the somatosensory cortex to quantify effects of scalp-to-sensor.

OPM and SQUID data were processed the same way using MNE-Python (Gramfort et al., 2013). The raw MEG data was high-pass filtered at 4 Hz, and homogeneous field compensation (HFC) was applied (Tierney et al., 2021). The annotation tool in MNE Python was used to omit stimulus artefacts and large transient deviations in the raw data. The SEF data was obtained by averaging trials for the time window of -100 to +50 ms around the electrical stimulus; the average signal over the pre-stimulus baseline of -100 to -5 ms defined the zero level in each sensor. The latency of the N20 response was identified using the runs with the smallest scalp-to-sensor distance for each sensor type, and the N20 magnitude (*B*_N20_) at that latency was then determined for all runs. The noise level was defined as the standard deviation (*σ*_N20_) of the unaveraged evoked response data across trials at the latency of the N20. SNR of the N20 peak was thus

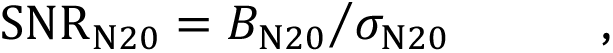

estimated for each run in all sensors. The spectral content of the continuous data was analyzed by calculating the power spectral density (PSD).

## 3. RESULTS

### 3.1. Simulations of SNR as a function of source depth

An example of the simulated signal *B*^max^ and the corresponding SNR in on-scalp (“OPM”) and off-scalp (“SQUID”) sensors as a function of the depth of a current dipole source is shown in **Fig. 3**. An adult-size spherical head model was assumed with relative noise level *η* = 3 (OPM on scalp, SQUID 18 mm off scalp, head radius 95 mm). Both signal and SNR decreased non-linearly with source depth. The signal was always stronger in the on-scalp sensor (**Fig. 3A**). The SNR curves, however, intersected at an intermediate dipole depth *d*_eq_ = 28 mm (**Fig. 3B**). For dipole locations more superficial than the equal-SNR depth (*d* < *d*_eq_) the SNR was higher for on-scalp than off-scalp sensors, whereas for deep locations (*d* > *d*_eq_) the opposite was found. This simulation illustrates that, although on-scalp sensors yield greater signal strength at any source depth, the SNR can be similar or even higher for off-scalp sensors when the sources are deep.

**Figure 3:**
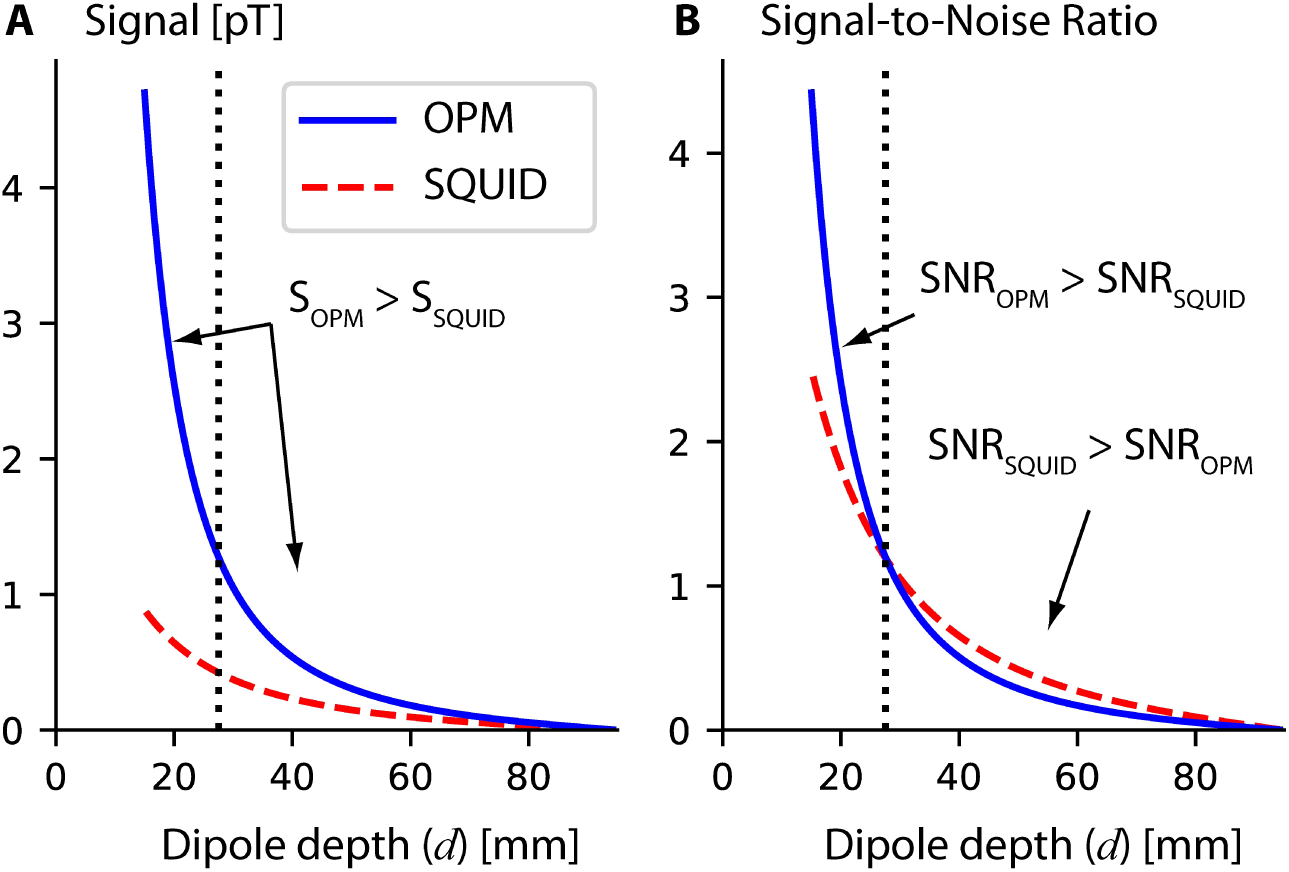
Simulated signal strength (**A**) and SNR (**B**) in on-scalp (OPM) and off-scalp (SQUID) MEG sensors measuring the radial component of the magnetic field due to a current dipole at different depths. The relative noise level was *η* = 3, the head radius ℎ = 95 mm, and the brain radius *b* = 80 mm. The source (dipole) depth *d* was measured from the head (scalp) surface. The scalp-to-sensor distance was ξ = 0 mm (on-scalp) for OPM and ξ = 18 mm (off-scalp) for SQUID sensors. An equal-SNR depth was found at *d*_eq_ = 28 mm from the scalp.

The effect of the relative noise level *η* between the on-scalp and off-scalp sensors on the SNR is illustrated in **Fig. 4**. Because the noise level in a given sensor was assumed to be independent of the source of interest, the shape of the depth-dependence curve for the SNR is the same as the shape for the signal amplitude, just scaled by the inverse of the noise level. However, because the shapes are different for on-scalp and off-scalp sensors, the SNR curves may intersect for certain values of *η*. When the noise levels were nearly equal (*η* = 1.1; **Fig. 4A**), the curves did not intersect: the SNR was higher for on-scalp sensors for all source depths (except for the sphere origin *d* = 95 mm, for which the signal vanished for all sensors). For *η* = 2.5 (**Fig. 4B**), the SNR curves intersected at the equal-SNR depth *d*_eq_ = 34 mm, such that the SNR was higher for on-scalp sensors only for sources more superficial than *d*_eq_. Increasing the relative noise level to *η* = 4.0 (**Fig. 4C**), decreased the value of *d*_eq_ to 19 mm, indicating that the SNR was higher for on-scalp sensors only for the most superficial sources. When the noise level was much higher in the on-scalp than off-scalp sensors (*η* = 6.0; **Fig. 4D**), no *d*_eq_ was found as the SNR was lower for on-scalp sensors for all source depths.

**Figure 4:**
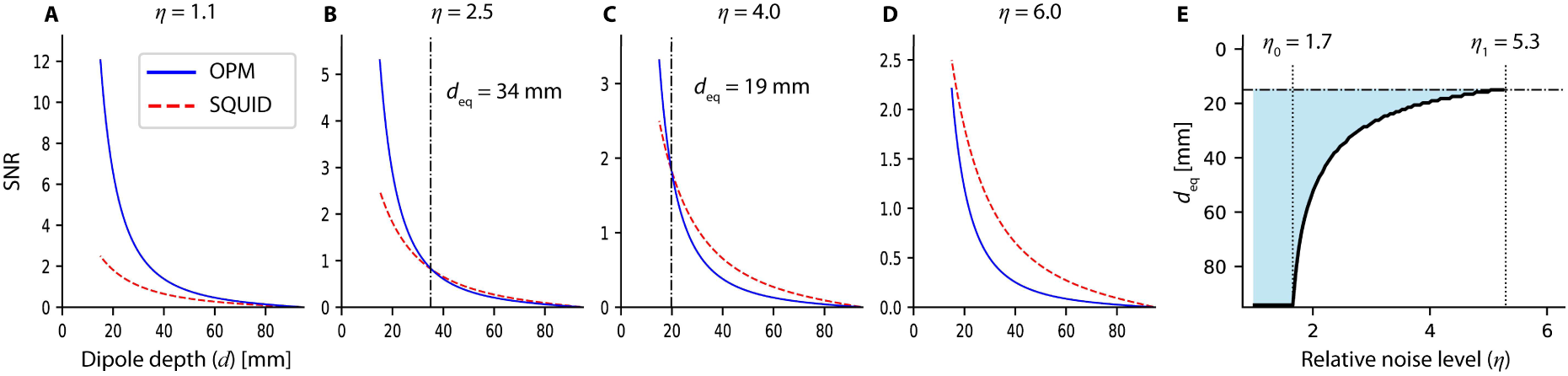
Effect of the relative noise level *η* on the simulated SNR in on-scalp (OPM) and off-scalp (SQUID) sensors. The head model and the scalp-to-sensor sensor distances as in Fig. 3. **A-D**: The SNR as a function of the dipole depth *d* (measured from the scalp surface) for four different relative noise levels: *η* = 1.1 (**A**), 2.5 (**B**), 4.0 (**C**), and 6.0 (**D**). The vertical dotted lines in **B** and **C** indicate the equal-SNR depth *d*_eq_. **E**: Equal-SNR depth *d*_eq_ as a function of *η*. The shaded area indicates the combinations of *d* and *η* values for which *d* < *d*_eq_ and correspondingly SNR_OPM_ > SNR_SQUID_.

The dependence of *d*_eq_ on the relative noise level *η* is summarized in **Fig. 4E**. For source locations above the curve (shaded area) SNR_OPM_ > SNR_SQUID_, whereas below the curve SNR_OPM_ < SNR_SQUID_. For near-equal noise levels (the leftmost part of the graph, *η* < 1.7), the SNR was higher in on-scalp sensors for all source depths. In the intermediate range (middle part, 1.7 < *η* < 5.3), the SNR in on-scalp sensors was higher than the SNR in off-scalp sensors only for the superficial source locations (*d* < *d*_eq_), whereas for deep sources (*d* > *d*_eq_) the SNR was higher in off-scalp than in on-scalp sensors. For very high on-scalp sensor noise (rightmost part, *η* > 5.3), the SNR was higher in off-scalp sensors for all source locations. Thus, the SNR in on-scalp sensors can be higher or lower than in off-scalp sensors, depending on the source location and the relative noise level.

The dependence of *d*_eq_ on *η* is shown for different head sizes in **Fig. 5**. For all head models, the dependence was qualitatively similar, with the equal-SNR source locations becoming more superficial (i.e., smaller *d*_eq_) when the noise level in on-scalp relative to off-scalp sensors got higher (**Fig. 5A**). Normalized *d*_eq_ values (relative to the brain radius *b*) suggest that for a given noise ratio *η*, there was a wider relative range of source depths for which SNR_OPM_ > SNR_SQUID_ (corresponding to the shaded area in **Fig. 4E**) in the smaller head models compared with the adult-size head model (**Fig. 5B**). For example, for *η* = 3, the normalized *d*_eq_ was 50% of the brain radius in the infant model, whereas it was only 15% in the adult model. This simulation suggests that the SNR benefit for superficial sources for on-scalp sensors is expected to apply to a larger proportion of the brain volume in smaller heads.

**Figure 5:**
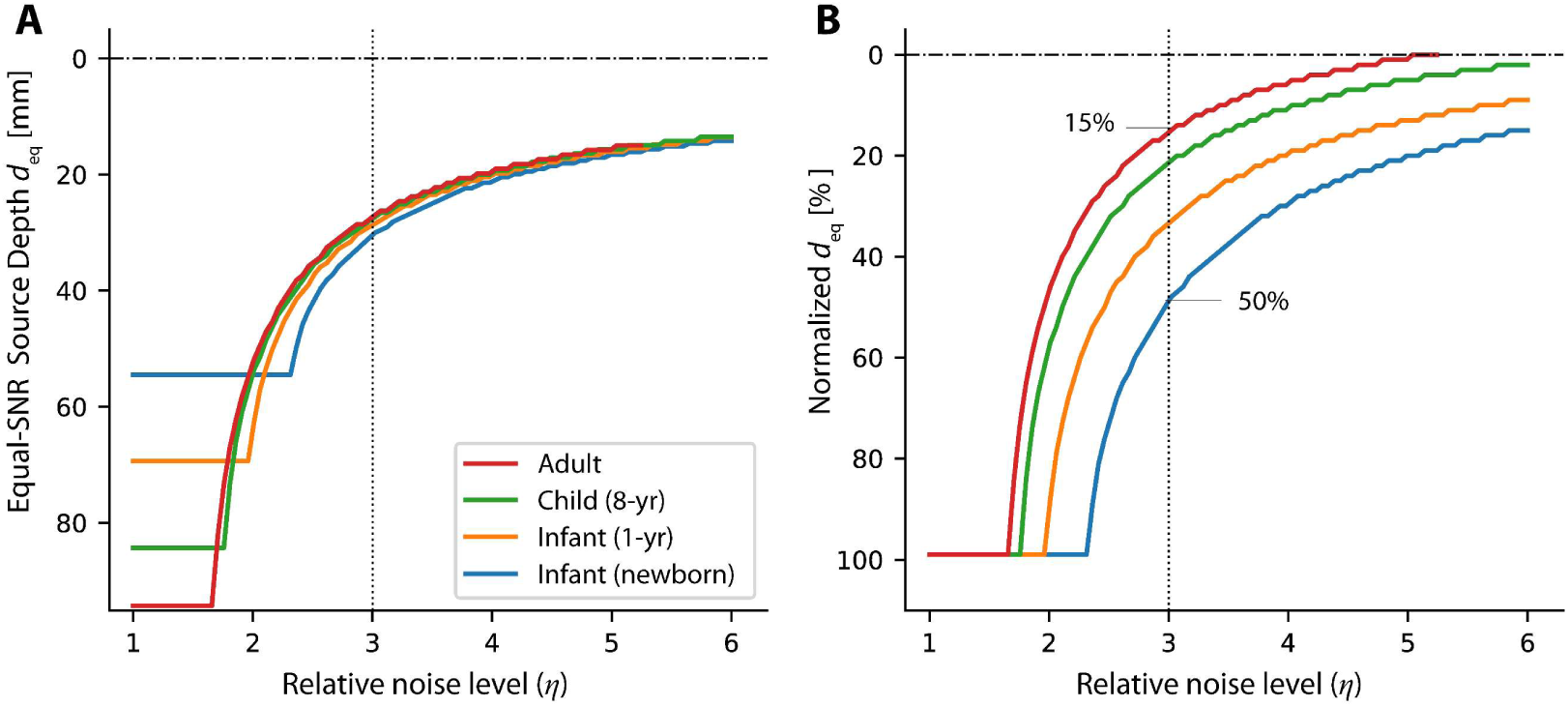
Simulated equal-SNR depth *d*_eq_ as a function of the relative noise level *η* for different head models, ranging from newborn to adult (see **Table 1**). In each case, the OPM sensor was assumed to be on-scalp (*∂* = 0 mm) and the SQUID off-scalp (*∂* = 18 mm). **A**: Absolute values of *d*_eq_. **B**: Normalized values of *d*_eq_ as percent of the brain radius. For a given relative noise level, the smaller head models showed larger normalized *d*_eq_ values (e.g., for *η* = 3, newborn: *d*_eq_ = 50%, adult: *d*_eq_ = 15%).

To relate the simulations more closely to our SEF experiment, we also varied the scalp-to-sensor distance ξ keeping the source location fixed. Here we assumed that both the signal and the noise depended on ξ, mimicking a situation in which the brain noise is the dominant noise source. Simulated *B_r_*^max^ (denoted by *B*_1_, *B*_2_, and *B*^3^) a function of ξ for 3 current dipoles located at different depths (*d*_1_ = 32 mm, *d*_2_ = 48 mm, and *d*_3_ = 64 in the adult-size head model) are shown as in **Fig. 6A**. As expected, the signals were higher when the sensors were closer to the scalp (right side of **Fig. 6A**) for each dipole location.

**Figure 6:**
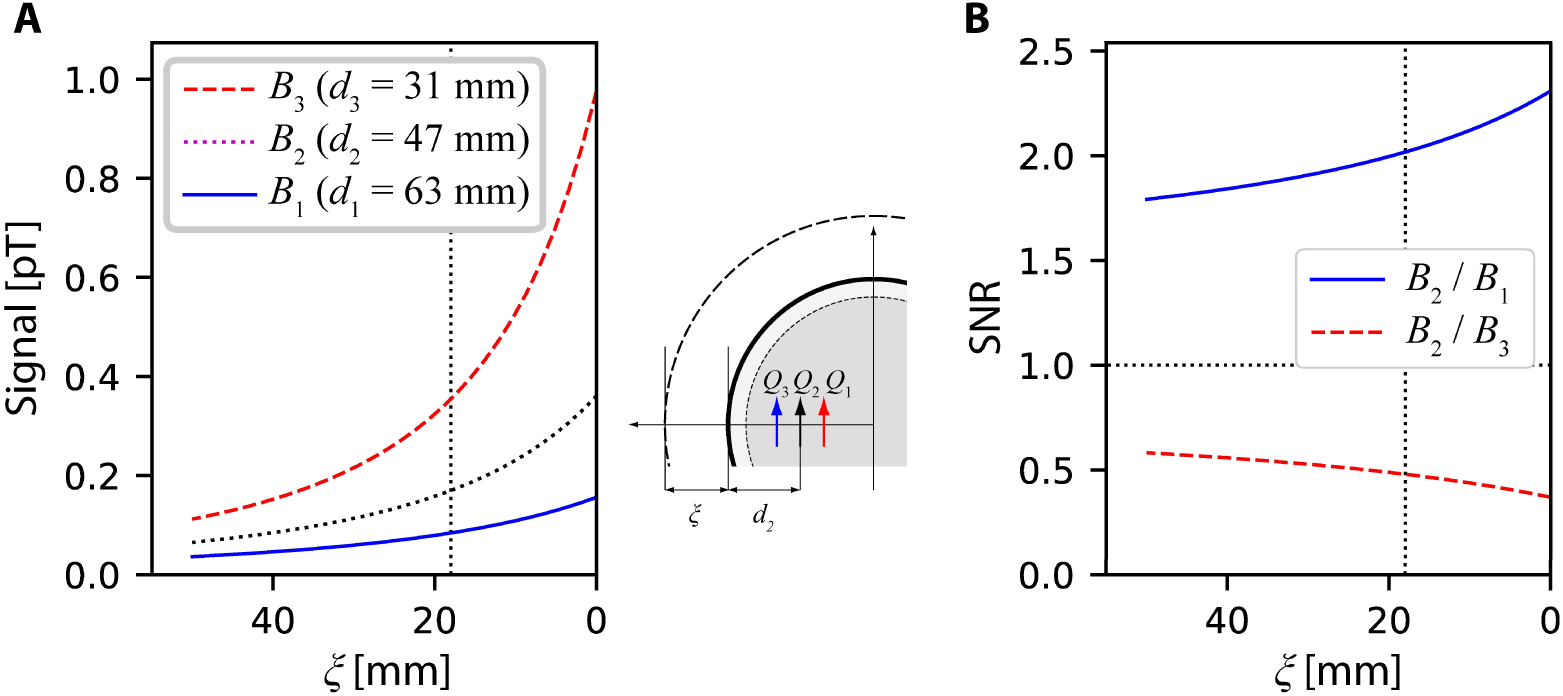
Dependence of the simulated signal and SNR on the scalp-to-sensor distance ξ. **A**: The magnetic field *B_r_*^max^ calculated for three different dipole depths (*B*_1_: *d*_1_ = 63 mm; *B*_2_: *d*_2_ = 47 mm; *B*_3_: *d*_3_ = 31 mm) in the adult-size head model (ℎ = 95 mm). The vertical line indicates the minimum scalp-to-sensor distance for the Triux SQUID system (ξ = 18 mm). **B**: The relative size of the magnetic field for the middle dipole location was compared with the field from the deeper (*B*_2_/*B*_1_) and the more superficial (*B*_2_/*B*_3_) sources

To simulate the dependence of the SNR on the scalp-to-sensor distance, we assumed that *B*_2_ was the signal of interest and the noise equaled either *B*_1_ or *B*_3_ (deep and superficial noise source, respectively). The corresponding SNR as a function of ξ are shown in **Fig. 6B**. When the noise source was deeper than the source of interest, the SNR (*B*_2_⁄*B*_1_, blue curve in **Fig. 6B**) was highest for the shortest scalp-to-sensor distance (i.e., on-scalp, ξ = 0 mm). In contrast, when the noise source was more superficial than the source of interest, the SNR (*B*_2_⁄*B*_3_, red dashed curve in **Fig. 6B**) was higher for off-scalp (non-zero ξ) than on-calp sensor placement. This can be seen as a consequence of the relative change in source-to-sensor distance being larger for the superficial noise source than for the source of interest as ξ varies. Even though a single dipole is not a realistic source model for brain noise, this simulation helps to illustrate qualitatively how SNR may depend on scalp-to-sensor distance when a noise source is located either deeper or more superficial than the source of interest.

### 3.2. Somatosensory evoked fields at different scalp-to-sensor distances

Averaged SEF responses in one OPM and one SQUID sensor at different scalp-to-sensor distances are shown in **Fig. 7A**. Overall, the N20 magnitude in OPM was about 3 times as large as in SQUID; however, the time courses were similar in the two types of sensors. The topographic maps showed characteristic dipolar patterns over the left somatosensory cortex (**Fig. 7B**).

**Figure 7:**
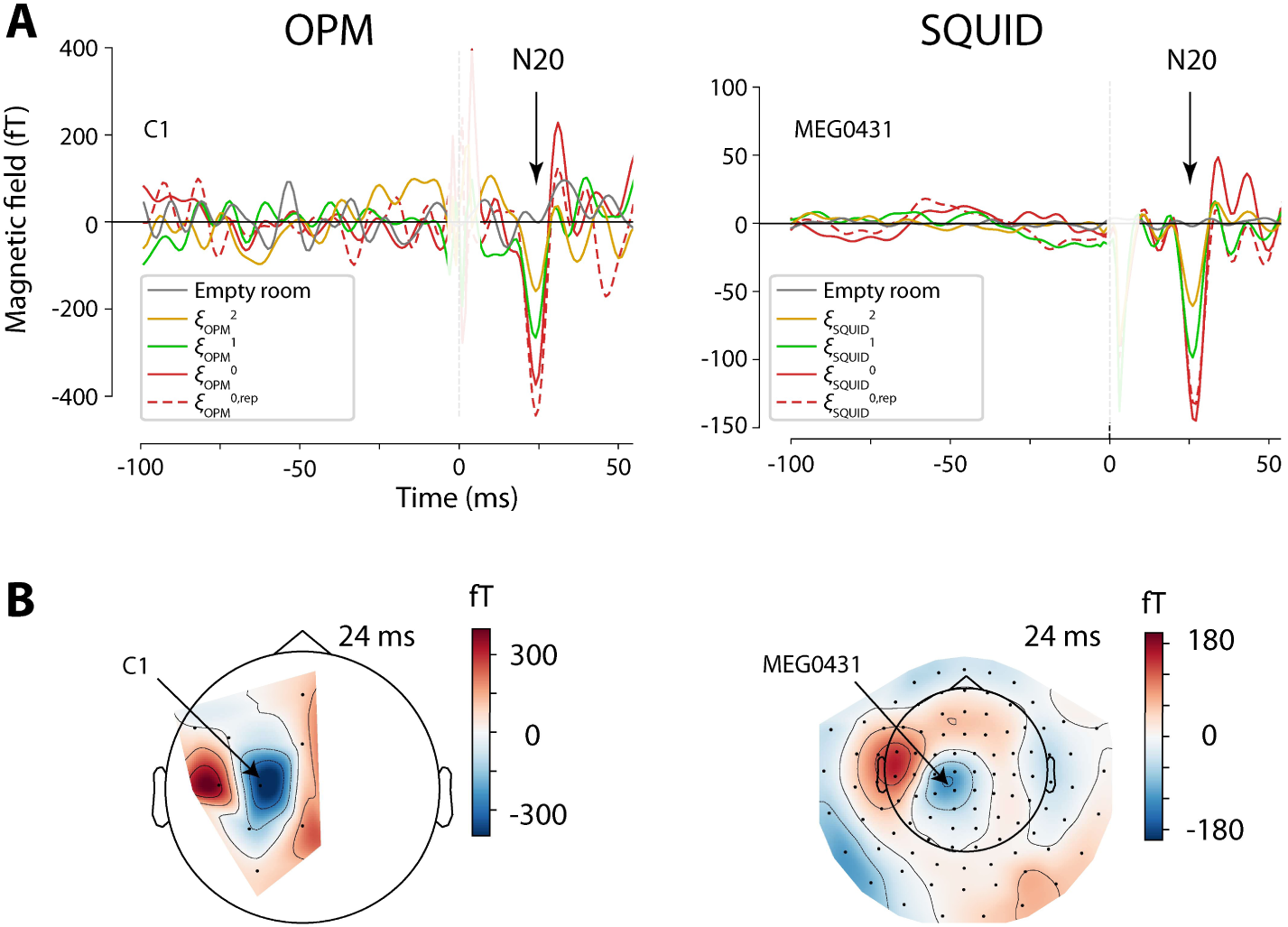
Somatosensory evoked magnetic fields. (**A**) Sensor time courses from 100 ms before to 50 ms after the electrical stimulation of the right median nerve at different OPM and SQUID sensor placements near the left somatosensory cortex. The placements closest to the scalp for OPM and SQUID are denoted by ξ^0^_OPM_ and ξ^0,rep^_OPM_, ξ^0^_SQUID_ and ξ^0,rep^_SQUID_; the others were about 1 and 2 cm further away from the scalp (ξ^1^_OPM_ and ξ^2^_OPM_, ξ^1^_SQUID_ and ξ^2^_SQUID_) Prominent N20 responses were seen in all runs. Empty room data was recorded without the participant. The stimulus artefacts near 0 ms are masked out to better visualize the evoked responses. Note that the vertical scale is different for OPM and SQUID. (**B**) Isocontour maps for the N20 peak latency for the closest-to-scalp placements (two runs averaged).

As expected, for both OPM and SQUID the N20 magnitude (*B*_N2_) was lower when the sensors were further away from the scalp (**Fig. 8A**). For SQUID, also the standard deviation *σ*_N2_ was lower when the sensors were further away from the scalp: *σ*_N20_(ξ^1^_SQUID_)/*σ*_N20_(ξ^0^_SQUID_) = 79%, *σ*_N20_(ξ^2^_SQUID_)/*σ*_N20_(ξ^0^_SQUID_) = 59% (**Fig. 8B**). This is consistent with the noise being dominated by brain noise, hence diminishing with distance from the brain. In contrast, for OPM, the *σ*_N20_ depended only weakly on the scalp-to-sensor distance: *σ*_N20_(ξ^1^_OPM_)/*σ*_N20_(ξ^0^_OPM_) = 91%, *σ*_N20_(ξ^2^_OPM_)/*σ*_N20_(ξ^0^_OPM_) = 101%, suggesting a smaller role for brain noise relative to environmental, instrumentation, or some other noise of non-brain origin. Because in SQUID both the signal and the noise, but in OPM mostly only the signal, were lower for larger scalp-to-sensor distances, the SNR depended less on the scalp-to-sensor distance in the SQUID than in the OPM data (**Fig. 8C**). For example, the SNR ratios in OPM were SNR_N20_(ξ^1^_OPM_)/SNR_N20_(ξ^0^_OPM_) = 68%, SNR_N20_(ξ^2^_OPM_)/SNR_N20_(ξ^0^_OPM_) = 31%, whereas the corresponding ratios in SQUID were 87% and 73%.

**Figure 8:**
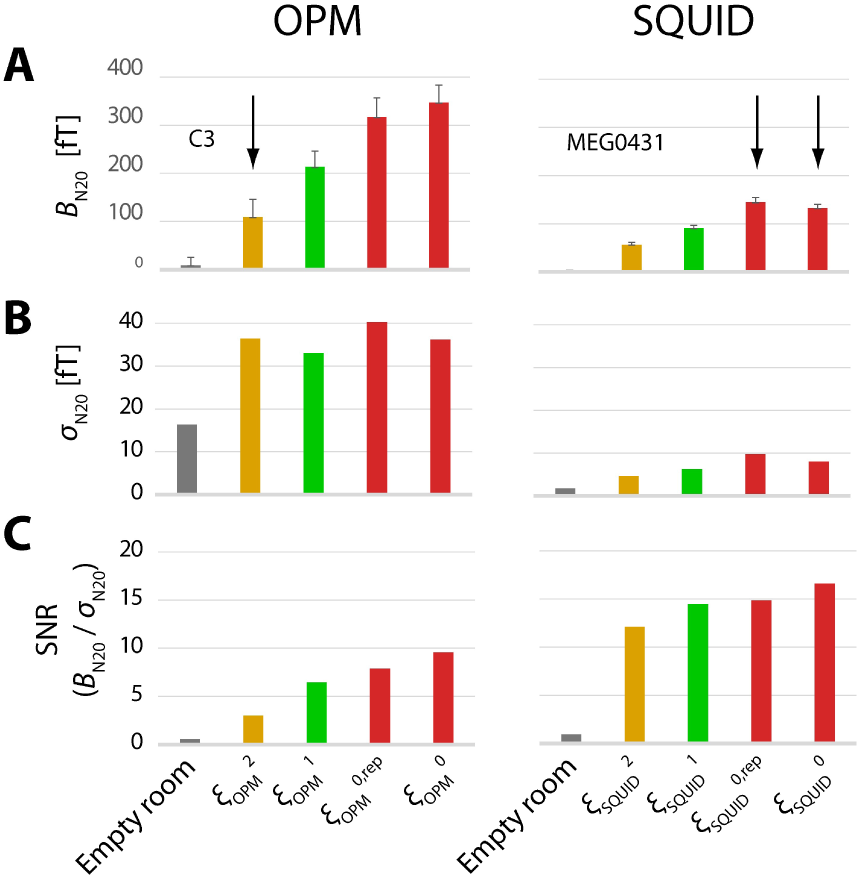
The SEF N20 response in OPM and SQUID at different distances from the scalp: (**A**) magnitude (*B*_N20_), (**B**) standard error of mean (σ_N2_) and (**C**) the corresponding signal-to-noise ratio (SNR). The arrows in **A** indicate cases the scalp-to-sensor distance was similar in OPM and SQUID (∼ 2 cm; ξ^2^*_OPM_*, ξ^0^*_SQUID_*).

Power spectrum densities of the raw data are shown in **Fig. G**. The PSDs for empty room data were relatively flat over a wide range of frequencies in both OPM and SQUID. At higher frequencies (above ∼35 Hz for OPM and ∼60 Hz for SQUID), PSD levels for all sensor placements were similar to those for empty-room data, suggesting that intrinsic sensor noise was dominant at these frequency ranges. At mid-frequencies (∼8–35 Hz for OPM, ∼4– 60 Hz for SQUID), the PSDs for both OPM and SQUID showed higher values when the sensors were closer to the scalp, suggesting that fields due to brain activity were dominant. In OPM, there was a large, possibly cardiac-related low-frequency component (∼4–8 Hz), which was independent of the scalp-to-sensor distance.

**Figure 9:**
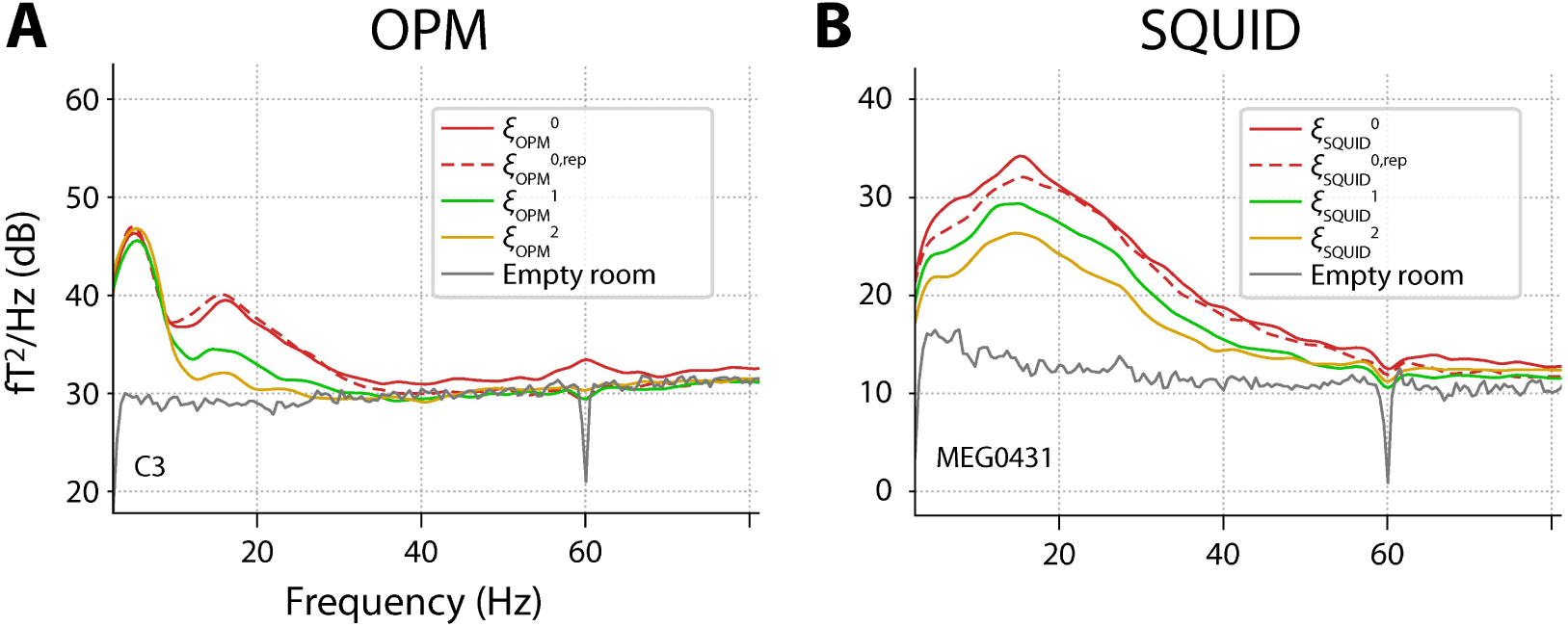
Power spectral density (PSD) of OPM (*left*) and SQUID (*right*) data at different sensor placements. Both sensors showed dependence on scalp-to-sensor distance in a mid-frequency range, peaking at ∼15 Hz.

## 4. DISCUSSION

The simulations suggested that there may exist an equal-SNR source depth for on-scalp vs. off-scalp MEG sensors. For sources located deeper than this depth, the SNR would be higher in off-scalp than on-scalp sensors. The equal-SNR depth depended on the relative noise level between on-scalp and off-scalp sensors. For a given relative noise level, the proportion of brain volume where the simulated SNR was higher in OPM than in SQUID was larger for child-than adult-sized head models. Somatosensory evoked responses recorded with OPM and SQUID sensors at varying distances from the scalp enabled us to identify whether the dominating contribution at different frequency ranges were of brain or non-brain origin, in general accordance with the stimulation findings and assumptions. The results help to better understand factors contributing to the relative SNR between on-scalp and off-scalp recordings of event-related data.

### 4.1. Sensitivity of MEG sensors to superficial vs. deep sources

Despite the substantially larger signal magnitude in on-scalp sensors, the SNR tended to be comparable between the two sensor types, as illustrated in **Fig. 3**. This dissociation between signal magnitude and SNR has been observed, e.g., for epileptic spikes and sensory- and motor-related responses (Brickwedde et al., 2024). The simulation results are consistent with the hypothesis that for event-related activity, the benefit gained from on-scalp placement of MEG sensors is largest for superficial sources in the brain (Boto et al., 2017; Iivanainen et al., 2017). For deep sources, off-scalp sensors can provide an SNR comparable to, or even higher than, that of on-scalp sensors. This may appear counter-intuitive considering that the on-scalp sensors are closer to all sources, including the deep ones. A combination of two contributions, however, influences the relative SNR: first, the relative difference in the signal magnitude for superficial vs. deep sources is larger for on-scalp than off-scalp sensors, and second, the brain noise is expected to be higher for on-scalp sensors.

Regarding the first contribution, faster drop in signal magnitude for on-scalp sensors as a function of source depth is evident, e.g., in **Fig. 4D**. (Note that the curves for SNR and signal magnitude have the same shape, they differ only by a constant factor, i.e., the noise level.) Enhanced sensitivity of on-scalp sensors to superficial sources has been demonstrated previously in simulations in which the dominant noise was assumed to be either intrinsic sensor noise (Boto et al., 2017; Iivanainen et al., 2020) or brain noise (Matsubara et al., 2026).

The reasons for the second contribution, i.e., higher noise level in on-scalp sensors, are two-fold. On the one hand, in the state-of-the-art sensor technology, the intrinsic sensor noise is higher in OPMs and high-T_c_ SQUIDs than in conventional liquid-helium cooled SQUIDs (Brookes et al., 2022). The intrinsic sensor noise level in conventional SQUID systems is so low that normally the background brain activity (brain noise) is the limiting factor for the sensitivity to transient events (M. Hämäläinen et al., 1993). On the other hand, the brain noise will be higher in on-scalp sensors due to the shorter distance between the sensors and the sources of the brain noise. Environmental noise is likely to affect on- and off-scalp sensors equally (however, cf. (Safar et al., 2024) for a discussion of artefacts due to implants that move relative to SQUIDs but not OPMs).

For the detectability of event-related activity, it is thus important to take into account realistic levels of brain noise (Goldenholz et al., 2009; Hunold, Funke, Eichardt, Stenroos, & Haueisen, 2016) in addition to the lead-field (forward-model)-based sensitivity to sources at different locations in the brain (Hillebrand & Barnes, 2002). For example, Roos et al. examined lead-field based sensitivity patterns, finding that OPMs measure stronger net signals from across the cerebellum compared with SQUIDs (Roos, Hamalainen, & Iivanainen, 2026). In contrast, in a different study, SNR-based results have suggested that the OPMs are beneficial mainly for the superficial regions of the cerebellum (Matsubara et al., 2026).

Pediatric studies are an important application of OPM-MEG (Edgar et al., 2026; Feys et al., 2022; Rhodes et al., 2024; T. P. L. Roberts, Birnbaum, Bloy, & Gaetz, 2025). Fixed adult-size SQUID helmets present a challenge in terms of scalp-to-sensor distances as well as movement of the head with respect to the sensors (Gaetz et al., 2008; Wehner, Hamalainen, Mody, & Ahlfors, 2008). Flexible placement of on-scalp sensors allows the sensor array to be adjusted to small head sizes. The present simulations of the equal-SNR source depth for different head models indicated that the SNR-benefit of on-scalp recordings may be largest for small heads (see **Fig. 5**). For a given relative sensor noise level, the percentage of the brain volume where SNR_OPM_ > SNR_SQUID_ was higher for small than large head models. This result suggests an additional potential benefit of the use of OPMs in pediatric studies.

### 4.2. Noise level at different scalp-to-sensor distances

Varying the scalp-to-sensor distances in the SEF experiment enabled us to infer whether the dominant spectral power at specific different frequency range was likely due to brain activity as opposed to being of non-brain origin, possibly instrumentation noise, environmental noise, or non-brain subject-related artefacts. In OPM, three qualitatively different frequency ranges were observed. The PSD in the middle frequency range (10–30 Hz) varied as a function of the scalp-to-sensor distance, thereby suggesting a main contribution from background brain activity. In contrast, for higher frequencies the PSD was identical with and without the participant, and therefore likely due to instrumentation and/or environmental noise. For the low-frequency end of the spectrum the OPM PSD was higher with the participant than in the empty room recording but depended only weakly on the scalp-to-sensor distance; this component may be related to the cardiac cycle.

### 4.3. Limitations and future directions

Several simplifying assumptions were made in the present stimulations, including a single current dipole source, a spherically symmetric head model, and magnetometer sensors detecting only the radial field component. In addition, we did not model the properties of the brain noise beyond the assumption that it diminishes with increasing scalp-to-sensor distance. Changing any of these assumptions would likely affect the equal-SNR crossover depth, though the main conclusions are not expected to depend critically on them, as discussed below.

Current dipoles in spherical geometry provides a useful approximation to realistic situations. The current dipole is commonly used to represent focal MEG source activity within a spatially restricted region in the brain (Scherg & Von Cramon, 1986). In addition, signals from extended and distributed sources can be expressed as the sum of signals from a set of current dipoles (Ahlfors et al., 2010; M. S. Hämäläinen & Ilmoniemi, 1994). The sphere model has been valuable for practical MEG forward modeling, especially when the sphere is chosen to match the local curvature of the skull near the source of interest (Huang, Mosher, & Leahy, 1999). For more realistic head geometries (Vorwerk et al., 2014), the concept of equal-SNR depth can be generalized by defining an equal-SNR surface that partitions the source space into regions where the SNR is higher for one type or sensor or the other. Simulated equal-SNR surfaces for the cerebellum can be seen in (Matsubara et al., 2026).

Most SQUID MEG systems measure approximately the magnetic field normal to the scalp surface, corresponding to the radial component *B*_r_ used in our simulations. Many OPM sensors measure simultaneously multiple field components (Brookes et al., 2021). The tangential and radial components have different spatial patterns over the scalp, but all magnetic field components diminish non-linearly with source-to-sensor distance, and therefore the SNR properties are expected to be qualitatively similar.

MEG systems commonly have gradiometric sensors that measure the difference in the magnetic field between two or more locations (Vrba & Robinson, 2001). Partial analogy can be made with the comparison of on-scalp vs. off-scalp sensors and magnetometer vs. gradiometer sensors, including the concept of equal-SNR source depth. Signal magnitude diminishes faster as a function of source depth for gradiometers than magnetometers, especially for planar gradiometers (Cuffin & Cohen, 1979). Qualitatively, the SNR curves in **Fig.** 4 could also represent a comparison between planar gradiometers (corresponding to the blue curves, labeled “OPM”) and magnetometers (red curves, “SQUID”), placed at the same scalp-to-sensor distance. Interestingly, realistic simulation of brain noise superimposed on epileptic discharges have suggested higher SNR for planar gradiometers than for magnetometers when the source is superficial, as expected, but no significant differences in SNR for deep sources (Hunold et al., 2016). This result of Hunold et al. is analogous to the case depicted in **Fig. 4A**.

Studies with larger OPM sensor arrays, more participants, anatomically more realistic head models, and advanced noise reduction and other signal processing techniques including beamformers (Boto et al., 2016) to dissociate signal and noise components, could help to further elucidate the properties and capabilities of the different types of MEG sensors. One consequence of the higher relative sensitivity of on-scalp sensors to superficial sources is that more detailed forward modeling, particularly of skull defects (Lau et al., 2016; Lew et al., 2021; Vorwerk et al., 2014), may be necessary, as conductivity deviations can appear as virtual sources in MEG data (Ahlfors, Lew, Hamalainen, Ilmoniemi, & Okada, 2026).

Of particular interest in future studies is characterizing how brain noise varies with scalp-to-sensor distance. In our simulations (**Fig. 6**) we assumed that the noise level depended on the scalp-to-sensor distance the same way as the field from a current dipole at a fixed depth. Even though this model lacks a realistic spatial distribution of the sources of brain noise, it illustrated how the scalp-to-sensor dependence of the brain noise could affect the SNR. The magnetic fields generated by background brain activity may vary substantially across specific frequency bands, subject populations, individual subjects, and the state of a subject (Marzetti et al., 2013). To evaluate the SNR in sensor arrays, it will be important to characterize the ongoing background brain activity (Bijma & de Munck, 2008; Engemann & Gramfort, 2015; Hunold et al., 2016). Identifying the properties of the sources of interest as well as the background brain activity, in principle, allows optimization of the SNR for MEG recordings, which is relevant to instrumentation design, subject setup, as well as experimental design in general.

### 4.4. Conclusions

The SEF data illustrated how larger signal magnitude does not necessarily lead to higher SNR, particularly if the noise is dominated by background brain activity. The simulation results support the view that on-scalp MEG offers its greatest SNR advantage for superficial cortical sources. Theoretical analysis of the equal-SNR source depth for a pair of on-scalp and off-scalp sensors suggested that the SNR benefit of on-scalp sensors may extend to larger proportions of the brain in children compared with adults. Varying scalp-to-sensor distance helped to dissociate different types of noise contributions in the experimental data. These results are expected to have implications for optimizing MEG sensor arrays and experimental designs.

## FUNDING

Supported by NIH R21NS140619, R01NS112183, P41EB030006, and S10OD030469. The content is solely the responsibility of the authors and does not necessarily represent the official views of National Institutes of Health.

## Notes

### Competing Interest Statement

The authors have declared no competing interest.

